# Age, Sex and Alzheimer’s disease: A longitudinal study of 3xTg-AD mice reveals sex-specific disease trajectories and inflammatory responses mirrored in postmortem brains from Alzheimer’s patients

**DOI:** 10.1101/2023.12.23.573209

**Authors:** Alicia J. Barber, Carmen L. del Genio, Anna Beth Swain, Elizabeth M. Pizzi, Sarah C. Watson, Vedant N. Tapiavala, George J. Zanazzi, Arti B. Gaur

## Abstract

**Background:** Aging and sex are major risk factors for developing late-onset Alzheimer’s disease. Compared to men, women are not only nearly twice as likely to develop Alzheimer’s, but they also experience worse neuropathological burden and cognitive decline despite living longer with the disease. It remains unclear how and when sex differences in biological aging emerge and contribute to Alzheimer’s disease pathogenesis. We hypothesized that these differences lead to distinct pathological and molecular Alzheimer’s disease signatures in males and females, which could be harnessed for therapeutic and biomarker development.

**Methods:** We aged male and female, 3xTg-AD and B6129 (WT) control mice across their respective lifespans while longitudinally collecting brain, liver, spleen, and plasma samples (n=3-8 mice per sex, strain, and age group). We performed histological analyses on all tissues and assessed neuropathological hallmarks of Alzheimer’s disease, markers of hepatic inflammation, as well as splenic mass and morphology. Additionally, we measured concentrations of cytokines, chemokines, and growth factors in the plasma. We conducted RNA sequencing (RNA-Seq) analysis on bulk brain tissue and examined differentially expressed genes (DEGs) between 3xTg-AD and WT samples and across ages in each sex. We also examined DEGs between clinical Alzheimer’s and control parahippocampal gyrus brain tissue samples from the Mount Sinai Brain Bank (MSBB) study in each sex.

**Results:** 3xTg-AD females significantly outlived 3xTg-AD males and exhibited progressive Alzheimer’s neuropathology, while 3xTg-AD males demonstrated progressive hepatic inflammation, splenomegaly, circulating inflammatory proteins, and next to no Alzheimer’s neuropathological hallmarks. Instead, 3xTg-AD males experienced an accelerated upregulation of immune-related gene expression in the brain relative to females, further suggesting distinct inflammatory disease trajectories between the sexes. Clinical investigations revealed that 3xTg-AD brain aging phenotypes are not an artifact of the animal model, and individuals with Alzheimer’s disease develop similar sex-specific alterations in canonical pathways related to neuronal signaling and immune function. Interestingly, we observed greater upregulation of complement-related gene expression, and lipopolysaccharide (LPS) was predicted as the top upstream regulator of DEGs in diseased males of both species.

**Conclusions:** Our data demonstrate that chronic inflammation and complement activation are associated with increased mortality, revealing that age-related changes in immune response act as a primary driver of sex differences in Alzheimer’s disease trajectories. We propose a model of disease pathogenesis in 3xTg-AD males in which aging and transgene-driven disease progression trigger an inflammatory response, mimicking the effects of LPS stimulation despite the absence of infection.

## Background

Alzheimer’s disease, the most common form of dementia, is a progressive neurodegenerative disorder characterized pathologically by amyloid beta (Aβ) plaques, neurofibrillary tangles (NFTs) and neuropil threads of hyperphosphorylated tau, gliosis and neuroinflammation, as well as neuronal and synaptic loss [1, 2]. These neuropathological impairments create a cyclic cascade of damage, spanning a multidecade disease continuum, eventually producing symptoms of memory loss, impaired judgment and logic, personality and behavioral changes, and disorientation. In the final stages of the disease, verbal and non-verbal communication, purposeful movement, and swallowing are greatly diminished, necessitating around-the-clock care [3]. More than 55 million people worldwide currently have dementia, and it is expected that without effective preventative or curative therapeutic innovation, this number will reach 139 million in the year 2050 [4].

The most significant risk factor for Alzheimer’s disease is age, as the likelihood of developing the disease doubles every 5 years after the age of 65 [5]. Sex differences in life expectancy contribute to a greater lifetime risk of dementia in women, as females outlive males by 4-7 years on average worldwide [6, 7]. Women with Alzheimer’s also have significantly longer life expectancies than men with the disease [8-10]. However, despite their prolonged survival, women exhibit increased global Alzheimer’s neuropathological burden and greater cognitive impairment [11-14]. As a result, women with Alzheimer’s experience higher disability-adjusted life years due to disease compared to men [15]. The longstanding phenomenon of women presenting with higher morbidity but lower mortality rates compared to men throughout the aging process, across diagnoses, is known as the “male-female health-survival paradox [16, 17].”

Factors contributing to sex differences in biological aging and longevity include, but are not limited to, genetics, mitochondrial function, cellular senescence, proteostasis, and inflammatory response [18]. Franceschi et al. coined the term “inflammaging” to describe low-grade chronic, systemic, inflammation in aging associated with increased proinflammatory cytokine production, which is a substantial risk factor for morbidity and mortality in older adults [19]. While both sexes experience aging-associated changes in immune function [20, 21], males have been shown to experience more unfavorable changes and a greater rate of inflammaging [22, 23]. Inflammaging likely plays a critical role in the pathogenesis of age-related diseases [24], including Alzheimer’s, influencing the manifestation of disease and survival time from diagnosis.

Despite the immense impact of age and sex on Alzheimer’s incidence and severity, many questions remain regarding how these leading risk factors contribute to the molecular mechanisms of Alzheimer’s disease. It is especially unclear how disease trajectories, from initiation through progression of Alzheimer’s, differ between males and females. Mouse modeling remains one of the only solutions to examine changes in organ systems throughout disease progression, yet few long-term studies investigating sex as a key variable exist. 3xTg-AD mice are a widely used transgenic model to study Alzheimer’s which carry *APP*, *MAPT*, and *PSEN1* gene mutations [25]. They are one of the only models to develop both Aβ plaques and NFTs, in addition to exhibiting stark sexual dimorphism [26-30]. Since the initial generation of the model, it has been described by multiple groups and confirmed by the donating investigator that male 3xTg-AD mice no longer exhibit plaque and tangle pathology [31-33], yet few studies have examined the molecular underpinnings of this development.

The goal of our study was to uncover drivers behind sex-specific Alzheimer’s disease phenotypes that surface with age by establishing a comprehensive time course of neural and peripheral abnormalities in the 3xTg-AD mouse model. Given that aging and inflammaging affect all body systems, we looked outside of the central nervous system, examining changes in the liver, spleen, and plasma in addition to the brain. We hypothesized that sex differences in the biological aging process result in distinct disease trajectories in males and females. To test our hypothesis, we conducted a multi-year study, collecting tissues and plasma from male and female 3xTg-AD and control mice at incremental ages across mature adult, middle age, and old age life stages. The experimental design allowed for parallel sex-specific characterization of progressive pathology and gene expression changes within “healthy” and “diseased” aging mice. Sex-specific transcriptomic profiles observed in aged 3xTg-AD mice were further substantiated in clinical Alzheimer’s data, revealing clear sex differences in neuroinflammation and synaptic impairment consistent between species.

## Methods

### Animals

#### Colony Maintenance

The 3xTg-AD mice utilized were previously described [25, 26] and were obtained from The Jackson Laboratory (B6;129-Tg(APPSwe,tauP301L)1Lfa *Psen1tm1Mpm*/Mmjax; MMRRC Strain #034830-JAX). Homozygous expression for the mutations in the *PSEN1*, *APP*, and *MAPT* genes were confirmed with TaqMan Real-Time PCR assays and the colony was maintained using 3xTg-AD mice breeding pairs. Wildtype controls were bred in-house - C57BL/6J (B6; Strain #000664) females and 129S1/SvImJ males (129S; Strain #002448) were crossed to produce B6129SF1/J mice. Mice were housed 3-5 per cage, kept on a 12-hr light/dark cycle, and allowed ad libitum access to food and water. All animal procedures were approved by Dartmouth College’s Institutional Animal Care and Use Committee (IACUC). Blood samples obtained from sentinel mice underwent quarterly and yearly serology testing with the Multiplexed Fluorometric ImmunoAssay at Charles River Research Animal Diagnostic Services to confirm the absence of pathogens (Table S1).

#### Statistical Analysis

Animal survival probabilities were compared between sexes and strains using the likelihood ratio test, Wald test, and log-rank test for Kaplan-Meier analyses. A Cox proportional hazards model was used to examine the association of sex and strain with mortality risk.

### Histology

#### Antibodies

We obtained beta-amyloid recombinant rabbit monoclonal antibody (H31L21, catalog #700254), phospho-tau (Ser202, Thr205) mouse monoclonal antibody (AT8, catalog #MN1020), and tau mouse monoclonal antibody (HT7, catalog #MN1000) from Thermo Fisher Scientific. We obtained CD45 rabbit monoclonal antibody (D3F8Q, catalog #70257) and F4/80 XP rabbit monoclonal antibody (D2S9R, catalog #70076 from Cell Signaling Technology.

#### Staining

Hemibrains, spleen, and liver tissue samples were collected from euthanized animals following perfusion with PBS and heparin. Samples were subsequently fixed in formalin for 24 hours and then transferred to 70% ethanol. Tissues were stored in hinged biopsy cassettes and embedded in paraffin. Immunohistochemistry slides were cut at 4um and air dried at room temperature before baking at 60°C for 30 minutes. Automated protocol performed on the Leica Bond Rx included paraffin dewax, antigen retrieval and staining. Heat-induced epitope retrieval using Bond Epitope Retrieval 2, pH9 (Leica Biosystems AR9961) was incubated at 100°C for 30 minutes. Primary antibody was applied and incubated for 15 minutes at room temperature. Primary antibody binding was detected and visualized using the Leica Bond Polymer Refine Detection Kit (Leica Biosystems DS9800) with DAB chromogen and hematoxylin counterstain.

#### Quantification

Slides were imaged with a 40x slide scanner and converted to digital files. QuPath software was utilized to quantify antibody staining. For Aβ staining, plaques were outlined with the QuPath wand tool, and plaque surface areas were measured and summated across the sagittal brain section. Total brain surface area, excluding the olfactory bulb and cerebellum due to variation in tissue integrity across samples, was measured. Total plaque surface area out of total brain surface area produced normalized Aβ plaque surface area values for each sample. For all other staining, QuPath automated positive cell detection analyzed the designated regions of interest, generating a number of positive detections per mm^2^ of annotation area. For each batch of samples, negative controls that underwent all histology protocol steps except for primary antibody application were quantified using the same methods, and resulting values were subtracted from stained samples to normalize for background staining and technical variation.

#### Statistical Analysis

Data were analyzed by ordinary one-way analysis of variance (ANOVA) corrected for multiple comparisons with Tukey’s test, using a single pooled variance, on GraphPad Prism (Version 10.1.1). P-values less than 0.05 were considered statistically significant.

### Luminex

#### Plasma collection

Blood was collected from living animals retro-orbitally or from recently euthanized animals through the inferior vena cava. Animals were anesthetized with isoflurane during retro-orbital bleeds. Blood samples were transferred into 1.5 mL Eppendorf tubes with edetic acid (EDTA) for anti-coagulation (4uL EDTA/∼200 uL blood). Tubes were immediately placed on ice and then centrifuged at 4°C for 10 minutes at 1000 rcf. The supernatant was transferred to a new tube and centrifuged at 4°C for 3 minutes at 14,000 rpm. Plasma was transferred to a new tube and stored at -80°C until use.

#### Assay

Inflammatory proteins were measured using the MILLIPLEX Mouse Cytokine/Chemokine Magnetic Bead Panel – Premixed 32 Plex – Immunology Multiplex Assay (EMD Millipore. Corporation, Billerica, MA). Calibration curves from recombinant cytokine standards were prepared with threefold dilution steps in the same matrix as the samples. High and low spikes (supernatants from stimulated PBMCs and dendritic cells) were included to determine cytokine recovery. Standards and spikes were measured in triplicate, samples were measured once, and blank values were subtracted from all readings. All assays were carried out directly in a 96-well filtration plate (Millipore, Billerica, MA) at room temperature and protected from light. Briefly, wells were pre-wet with 100 uL PBS containing 1% BSA, then beads together with a standard, sample, spikes, or blank were added in a final volume of 100 uL and incubated together at room temperature for 30 minutes with continuous shaking. Beads were washed three times with 100 uL PBS containing 1% BSA and 0.05% Tween 20. A cocktail of biotinylated antibodies (50 uL/well) was added to beads for a further 30-minute incubation with continuous shaking. Beads were washed three times, then streptavidin-PE was added for 10 minutes. Beads were again washed three times and resuspended in 125 uL of PBS containing 1% BSA and 0.05% Tween 20. The fluorescence intensity of the beads was measured using the Bio-Plex array reader. Bio-Plex Manager software with five-parametric-curve fitting was used for data analysis.

#### Statistical Analysis

Data points considered out of range below (OOR<) were substituted with a 0, the minimum detectable value. Data were analyzed by ordinary one-way ANOVA corrected for multiple comparisons with Tukey’s test, using a single pooled variance, on GraphPad Prism (Version 10.1.1). P-values less than 0.05 were considered statistically significant.

### RNA Isolation and Sequencing

#### Isolation

Brain tissue was dissected from animals following perfusion with PBS and heparin. Samples were flash-frozen in liquid nitrogen and stored at -80°C. Samples were later placed in a deep-well 96-well plate with stainless steel RNase-free homogenization beads and QIAzol lysis buffer. Plates were inserted into the SPEX SamplePrep 1600 MiniG tissue homogenizer and run at maximum speed for 5 minutes. Following homogenization, the tissue solution was added to QIAzol lysis buffer and stored at -80°C. Lysed tissue solution was subsequently added to the QIAGEN miRNeasy Micro Kit for purification of total RNA.

#### Sequencing

RNA for RNA-seq was quantified by qubit, and integrity was measured on a fragment analyzer (Agilent). 200 ng RNA was hybridized to FastSelect probes (QIAGEN) for ribodepletion, followed by library preparation using the Kapa RNA HyperPrep kit (Roche) following the manufacturer’s instructions. Libraries were pooled for sequencing on a NextSeq500 instrument (Illumina) targeting 30M, paired-end 50bp reads per sample. Raw data was pre-processed by removing adapter sequences (cutadapt), aligning to the mm10 reference genome using HISAT2 and quantifying read counts using the “featurecounts” function from the subreads package to generate a gene expression matrix for downstream processing.

### RNA-Seq Analysis

#### Batch Adjustment

To adjust for sequencing batch effects within our total sample population, ComBat-seq was applied as previously described [34].

#### Differential Expression Analysis

DESeq2 was utilized as previously described [35] to identify differentially expressed mRNAs (DEGs) between 3xTg-AD and WT controls at various ages using R (Version 4.3.0) and RStudio (Version 2023.12.0+369). A negative binomial generalized linear model was applied. Differential expression p-values were adjusted for multiple comparisons with the Benjamini-Hochberg method.

#### Pathway Analysis

Statistically significant, differentially expressed genes (p<0.05) were categorized into canonical pathways using QIAGEN Ingenuity Pathway Analysis (IPA) Core Analysis [36] and Enrichr [37]. Predicted upstream regulators of DEGs were identified using IPA Upstream Analysis.

#### Over-Representation Analysis

Statistically significant, differentially expressed genes (p<0.05) underwent Over-Representation Analysis (ORA) to determine if specific gene ontology biological processes were likely present more than by chance. WebGestalt [38] was utilized to perform ORA with the Illumina mouseref 8 reference set. Affinity propagation was applied for redundancy reduction of enrichment categories. Differential expression p-values were adjusted for multiple comparisons by the Benjamini-Hochberg method.

## Results

### Multi-year longitudinal study of 3xTg-AD mice reveals sex-specific differences in life expectancy

We studied male and female 3xTg-AD mice across their lifespan and used B6129 (WT) mice as sex and age-matched controls. Brain, liver, and spleen tissues, in addition to plasma samples, were collected at successive time points starting at 3 months of age, with primary analyses being run on samples from mice aged 6-, 12-, 15-, 18-, and 21-months (n=3-8 mice of each sex and strain per age group). Survival was tracked to evaluate variances in longevity between groups (Fig. 1a). By the end of the study period, animals had either been sacrificed in good health for timepoint collection, sacrificed unexpectedly due to signs of imminent death (labored breathing, hunched posture, minimal movement, squinted eyes, frailty, etc.), found dead, or were still living. Data points from animals that were sacrificed for planned timepoints or still living, were censored in the survival analysis. Abnormalities observed in various organ systems during necropsy were described (Table S2), however, it is unknown if manifestations were directly related to disease progression, and most animals found dead were not dissected due to their state of decomposition.

**Figure 1.**
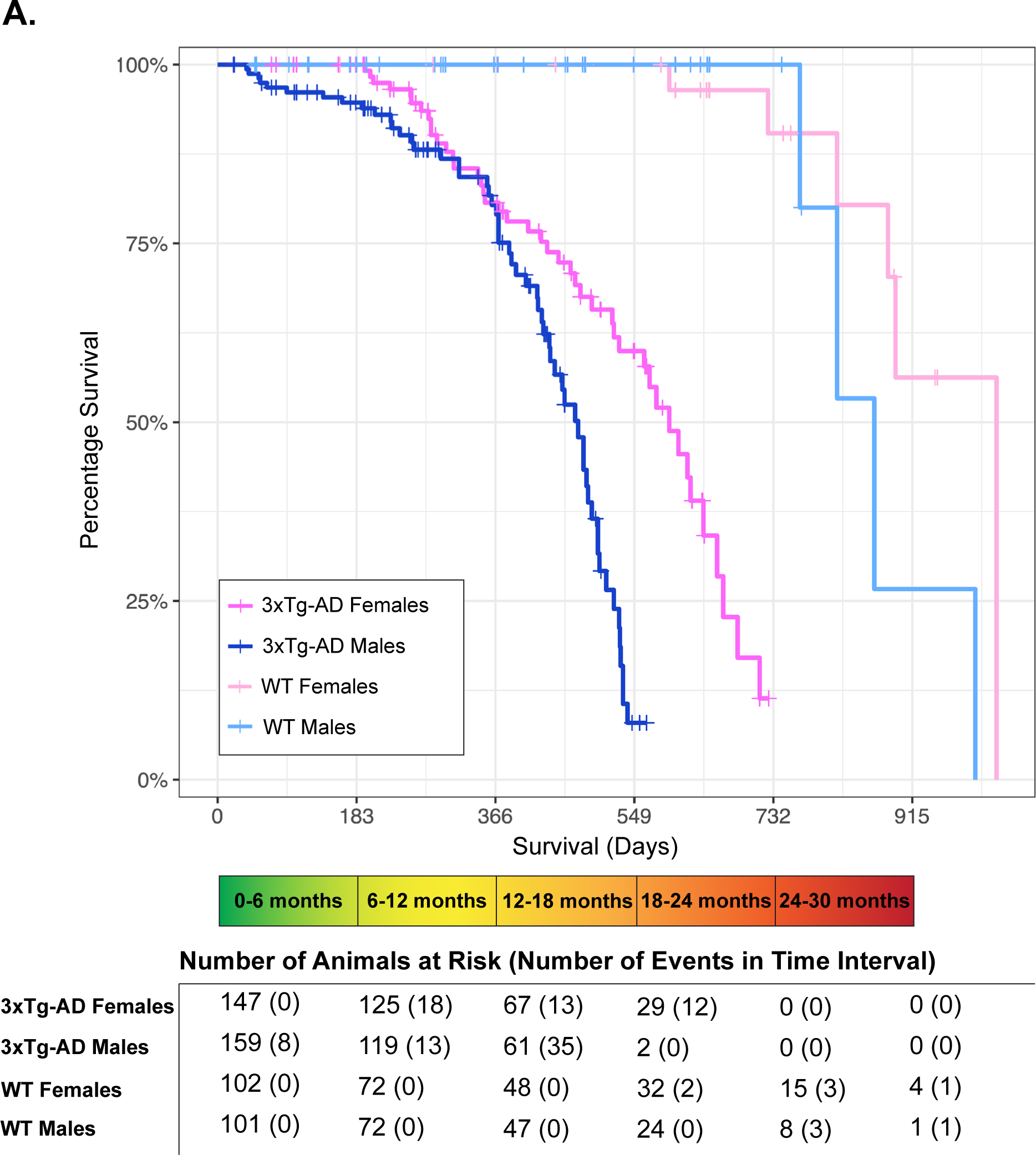
3xTg-AD females outlive males. (A) Kaplan-Meier curves for the survival of animals by sex and strain (3xTg-AD and B6129 (WT)). Data were available for 509 mice. Planned mortality events for timepoint sample collection were censored (+). Uncensored mortality events included animals that were found dead or sacrificed due to observed signs of imminent death. The number of animals at risk and the number of uncensored mortality events per 6-month time interval are shown below the graph.

3xTg-AD mutations were associated with a substantially increased risk of mortality, as 3xTg-AD mice had nearly 100 times the risk of mortality than controls (hazard ratio 95.2, 95% confidence interval 22.33-405.71, Table 1). WT mice experienced no natural mortality events up to 2-years of age (730 days), with animals surviving past 30 months. Compared to their respective controls, male 3xTg-AD life expectancy was 12 months shorter on average, while female 3xTg-AD life expectancy was 6 months shorter on average.

**Table 1.** Multivariate Cox Regression Analysis of Animal Survival.

| Characteristic | Hazard Ratio | 95% Confidence Interval | p-value |
| --- | --- | --- | --- |
| Sex (male) | 2.75 | 1.79-4.22 | 3.81E-16 |
| Strain (3xTg-AD) | 95.2 | 22.33-405.71 | 7.34E-10 |

The likelihood ratio test, Wald test, and log-rank test were all significant ( $p < 2E-16$ ). $n = 509$ mice, number of events = 109.

3xTg-AD male mice largely did not survive past 18 months of age, while 3xTg-AD females exhibited lifespans up to roughly 24 months. Male sex was associated with an increased risk of mortality, with a hazard ratio of 2.75 (95% confidence interval 1.79-4.22, Table 1). Overall, our results demonstrate that the 3xTg-AD mice have a significantly shortened lifespan, which is further exacerbated in male mice.

### Sex-specific severity of hallmark Alzheimer’s neuropathology

To characterize the initiation and progression of Alzheimer’s-related neuropathology, longitudinal immunohistochemistry analyses were run on sagittally-cut brain hemispheres in female and male 6-, 12-, 15-, 18-, and 21-month-old (oldest age dependent upon survival) (n=3-8 mice of each sex per age group) 3xTg-AD mice. Characteristic hallmarks of Alzheimer’s neuropathology, Aβ plaques, NFTs, and total tau expression were analyzed. Aβ antibody (H31L2) staining of brain samples was used to evaluate plaque accumulation throughout the course of the disease. We observed that extracellular Aβ plaques were first detectable at 12-months-old in the hippocampal formation of female 3xTg-AD mice, and plaque accumulation progressed significantly with age (Fig. 2a). By 15 months of age, plaques were also observed in the female 3xTg-AD pons, anterior olfactory nucleus, and cerebral cortex. Extracellular Aβ plaques were not detected in any male 3xTg-AD brain samples across the lifespan.

**Figure 2.**
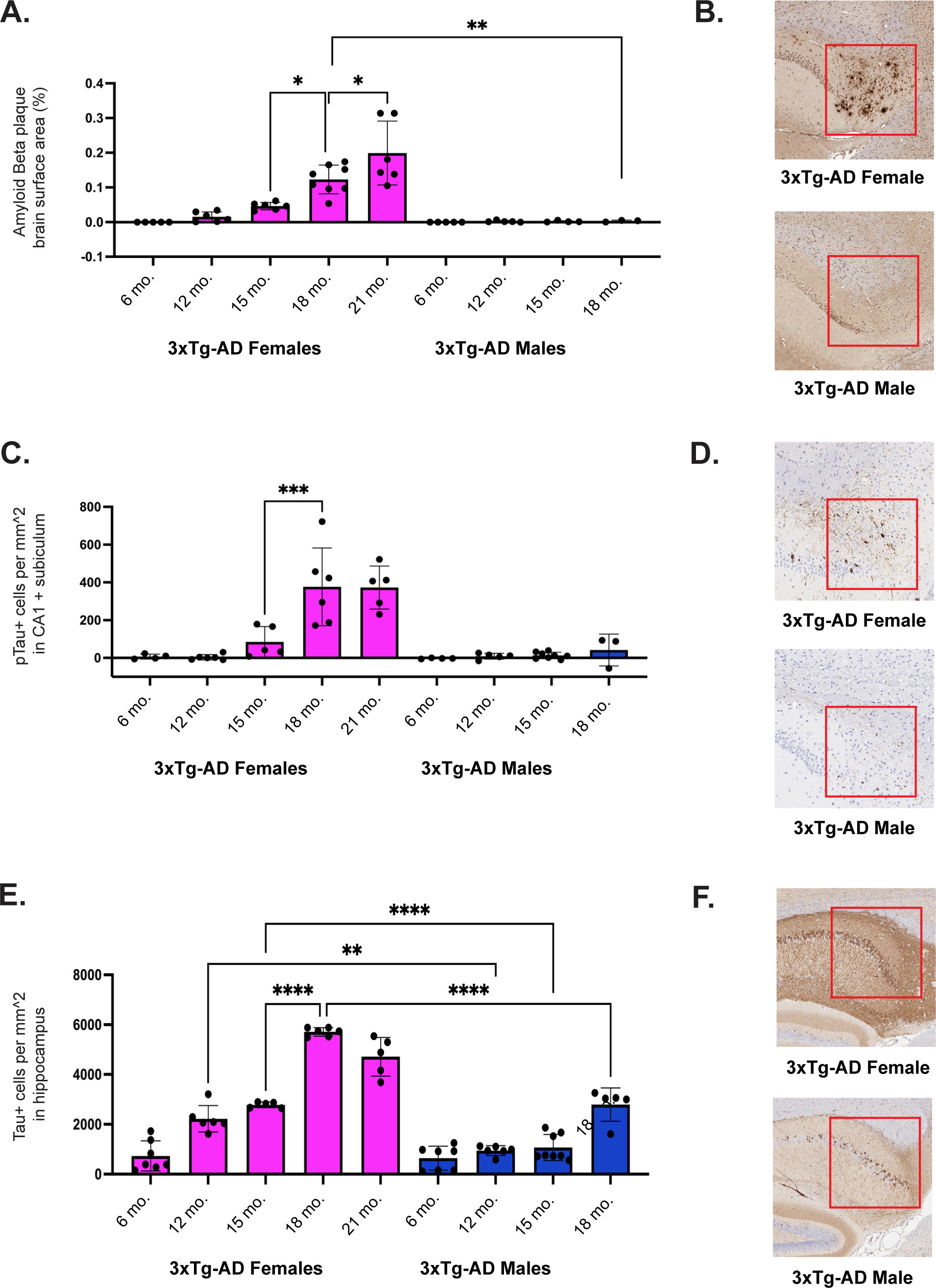
3xTg-AD females exhibit progressive Alzheimer’s disease neuropathology. (A) Quantitative analysis of Aβ plaque deposition across sagittal brain sections in female and male 3xTg-AD mice at 6-, 12-, 15-, 18-, and 21-months-old. (B) Representative images of hippocampal sections from 18-month-old female and male 3xTg-AD mice stained with a beta-amyloid-specific antibody (n=3-8/sex+age). Red squares highlight regions with differences in immunoreactivity between sexes. (C) Quantitative analysis of phosphorylated-tau (Ser202, Thr 205) positivity in the CA1 and subiculum of female and male 3xTg-AD mice at 6-, 12-, 15-, 18-, and 21-months-old. (D) Representative images of hippocampal sections from 18-month-old female and male 3xTg-AD mice stained with a phosphorylated-tau-specific antibody (n=3-8/sex+age). Red squares highlight regions with differences in immunoreactivity between sexes. (E) Quantitative analysis of total tau positivity surrounding the hippocampus of female and male 3xTg-AD mice at 6-, 12-, 15-, 18-, and 21-months-old. (F) Representative images of hippocampal sections from 18-month-old female and male 3xTg-AD mice stained with a tau-specific antibody (n=5-8/sex+age). Red squares highlight regions with differences in immunoreactivity between sexes. Data were analyzed by one-way ANOVA with multiple comparisons. Asterisks indicate significant differences between compared groups (* = p<0.05, ** = p≤0.01, *** = p≤0.001, **** = p≤0.0001). Error bars represent mean with SD.

Representative images of 18-month-old female and male 3xTg-AD hippocampal sections revealed a cluster of Aβ plaques in the female subiculum; however, the male brain does not present with plaque deposition (Fig. 2b). These results indicate that female 3xTg-AD mice exhibit an age-dependent accumulation of Aβ plaques. Further, Aβ protein aggregation is specific to females in the 3xTg-AD model, and plaque pathology is not contributing to the shortened lifespan of 3xTg-AD males.

To assess the accumulation of NFTs during the course of the disease, sagittally cut brain hemispheres were collected from female and male 6-, 12-, 15-, 18-, and 21-month-old 3xTg-AD mice (oldest age dependent upon survival) (n=3-8 mice of each sex per age group) and stained with a phospho-tau (Ser202, Thr205) antibody (AT8). NFTs and neuropil threads were quantified within the CA1 and subiculum. We observed that phospho-tau pathology started in 15-month-old 3xTg-AD female mice and progressed significantly with age (Fig. 2c). Slight tangle pathology was observed in 3xTg-AD male mice, with very minimal levels of accumulation measured at 18 months of age. Representative images of 18-month-old female and male 3xTg-AD hippocampal sections reveal a significantly higher density of tangles in the female subiculum compared to the male subiculum (Fig. 2d). Comparable analyses were conducted with a tau antibody (HT7) to measure levels of total tau expression (n=5-8 mice of each sex per age group). Tau-positive cells were quantified within a chosen region including and surrounding the hippocampal formation. We found that tau expression progressed with age, more significantly in 3xTg-AD females compared to males (Fig. 2e). Representative images of 18-month-old female and male 3xTg-AD hippocampal sections revealed a greater density of tau-positive cells in the female entorhinal cortex compared to the male entorhinal cortex. These results indicate that phospho-tau and total tau are significantly higher in the brains of 3xTg-AD female mice compared to males.

### Male 3xTg-AD mice demonstrate severe splenic and hepatic inflammation

To investigate the impact of disease progression outside of the central nervous system, we assessed spleen and liver histology. We measured splenic weight in 6- and 18-month-old, male and female, 3xTg-AD and WT mice (n=4-6 mice of each sex and strain per age group). 3xTg-AD male mice exhibited significantly enlarged spleens, with an average weight of 4.6 grams in 18-month-old 3xTg-AD male mice, 36x the size of age-matched control spleens, and nearly 15% of their total body weight (Fig. 3a). Additionally, hematoxylin and eosin staining revealed histological changes in 3xTg-AD male splenic tissue, with a relative decrease in white pulp. As splenomegaly developed, the pink vascular spaces of the spleen and lymphoid follicles were encroached upon by infiltrating neutrophils [39]. This results in the obstruction of venous outflow of blood and abnormal enlargement in size. Representative images of spleens from 18-month-old, female and male, 3xTg-AD and WT mice (Fig. 3b) captured these changes. Overall, these results suggest that 3xTg-AD male mice experience immense splenomegaly, possibly as a result of peripheral organ inflammation, at the end of their lives.

**Figure 3.**
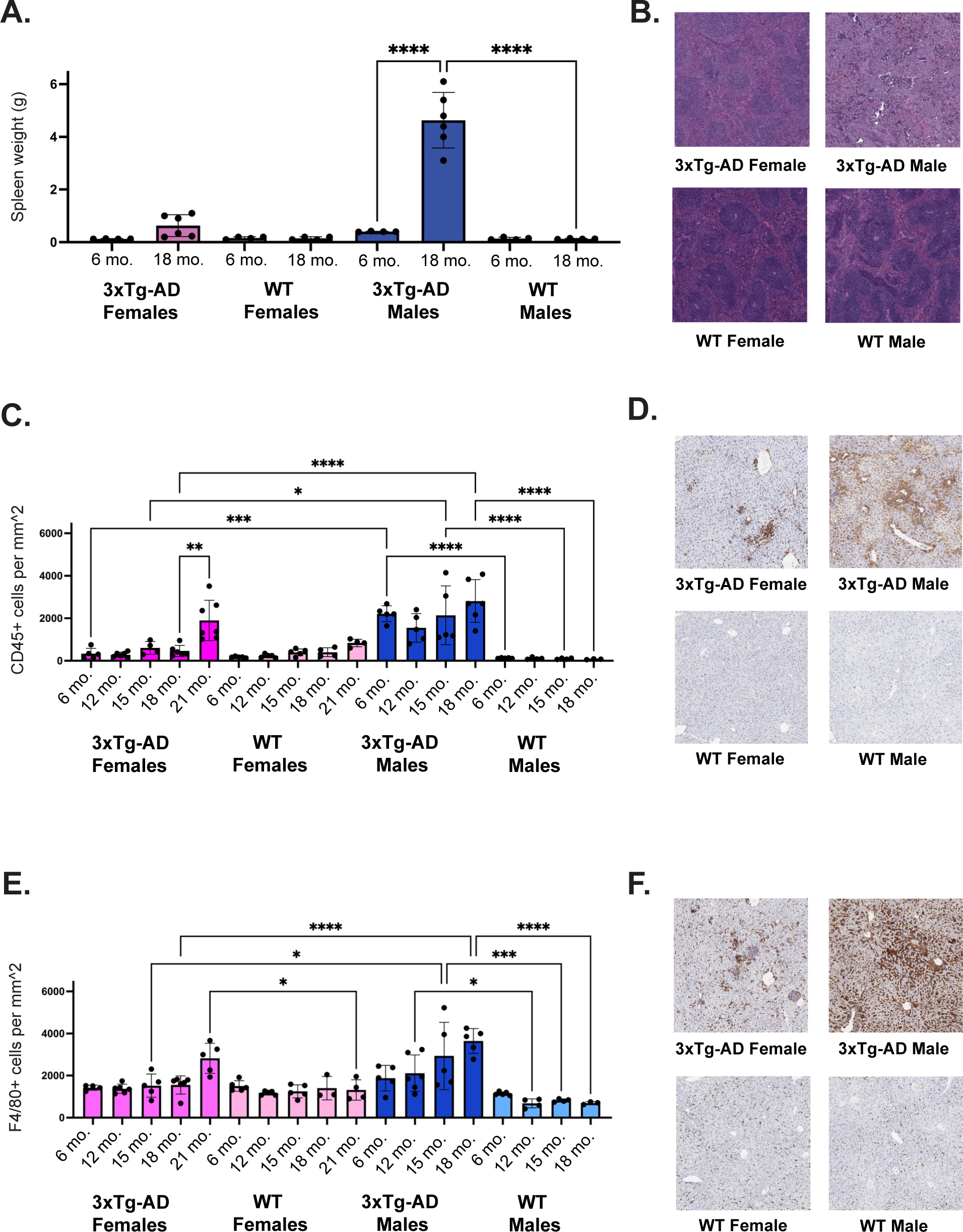
3xTg-AD males experience exacerbated age-dependent peripheral inflammation. (A) Splenic weight of female and male, 3xTg-AD and WT mice at 6-, and 18-months-old (n=4-6/sex+age). (B) Representative hematoxylin and eosin staining of spleen tissue in 18-month-old female and male, 3xTg-AD and WT mice depicting differences in white pulp organization. (C) Quantitative analysis of CD45 positivity in liver tissue from female and male 3xTg-AD mice at 6-, 12-, 15-, 18-, and 21-months-old. (D) Representative images of liver tissue from 18-month-old female and male, 3xTg-AD and wildtype mice, stained with a CD45-specific antibody (n=3-7/sex+age). (E) Quantitative analysis of F4/80 positivity in liver tissue from female and male, 3xTg-AD and WT mice at 6-, 12-, 15-, 18-, and 21-months-old. (F) Representative images of liver tissue from 18-month-old female and male, 3xTg-AD and WT mice, stained with an F4/80- specific antibody (n=3-6/sex+age). Data were analyzed by one-way ANOVA with multiple comparisons. Asterisks indicate significant differences between compared groups (* = p<0.05, ** = p≤0.01, *** = p≤0.001, **** = p≤0.0001). Error bars represent mean with SD.

To elucidate a potential source of peripheral organ dysfunction that could be contributing to splenomegaly, we measured two markers of inflammation in liver tissue throughout the course of disease progression. Liver tissue was collected from 6-, 12-, 15-, 18-, and 21-month-old male and female, 3xTg-AD and WT mice (oldest age dependent upon survival) (n=3-5 mice of each sex and strain per age group). Liver tissue was stained with a CD45 antibody to evaluate tissue wide inflammation. CD45 antigen, or leukocyte common antigen, is expressed on almost all hematopoietic cells except for mature erythrocytes [40]. We found that CD45 levels were significantly elevated in 3xTg-AD males compared to controls, as early as 6 months of age, while levels were not significantly elevated in 3xTg-AD females compared to controls until 21 months of age (Fig. 3c,d). To detect inflammation specifically resulting from increased macrophage infiltration, liver tissue was stained with F4/80, a marker for Kupffer cells - liver-specific macrophages. F4/80 levels were significantly elevated in 3xTg-AD males compared to controls, beginning at 15 months of age (Fig. 3e). However, in 3xTg-AD females a significant increase in macrophage infiltrates compared to controls was not observed until 21 months of age (Fig. 3e,f). Overall, these results indicate that 3xTg-AD males exhibit prolonged hepatic inflammation, which is present at 6 months of age and persists throughout the lifespan, likely contributing to splenomegaly and heightened mortality.

### Aged 3xTg-AD mice exhibit sex-specific plasma cytokine profiles

To investigate the systemic impact of aging and disease progression, we measured the concentrations of cytokines, chemokines, and growth factors in the plasma of male and female, 3xTg-AD and control mice, at 6-, 15-, 18-, 21-, and 24-months-old (oldest age dependent upon survival) (n=3-6 mice of each sex and strain per age group). We found that levels of interleukin (IL)-12, IL-10, IL-6, CXC chemokine ligand (CXCL) 2, and CC chemokine ligand (CCL) 4, were significantly increased in 3xTg-AD males only, between 15 and 18 months of age, while CXCL9 and CXCL10 levels were significantly increased in 3xTg-AD females only, at 24 months of age (Sup Fig. 1). Additionally, granulocyte-colony stimulating factor (G-CSF) and CXCL1 levels were greater in 3xTg-AD mice. G-CSF levels trended upward with age in both sexes but increased earlier in 3xTg-AD males. Overall, these results indicate that 3xTg-AD males exhibit exacerbated systemic inflammation relative to 3xTg-AD females, with more inflammatory proteins circulating in their bloodstream in late-stage disease progression and old age.

### 3xTg-AD mice manifest sex-specific timelines of immune-related gene expression in the brain

Transcriptomic profiling of female and male 3xTg-AD and WT brains was carried out across their lifespans to investigate potential molecular changes in the brain associated with the described neuropathology and systemic inflammation observed during the course of disease progression. RNA-Seq was performed on bulk brain tissue from animals between the ages of 6 and 24 months old, depending on lifespan (n=3-4 mice of each sex and strain per age group). Mice are considered mature adults from about 3-6 months of age. However, at about 3-months-old mice are just finishing a period of puberty, rapid aging, and maturational growth and are equivalent to about 20-years-old in human years. At 6-months-old, mice are equivalent to about 30-years-old in human years, and will begin experiencing age-related changes past this point [41, 42]. For these reasons, 6 months old was chosen as the mature adult baseline for comparisons in differential gene expression. A negative binomial generalized linear model was utilized to measure genes differentially expressed at progressive ages compared to a baseline of 6-months-old (Sup Fig. 2). We found that in 3xTg-AD females and males, numerous genes were increasingly dysregulated with age, exhibiting peak differential expression at the oldest time points (Fig. 4a,b).

**Figure 4.**
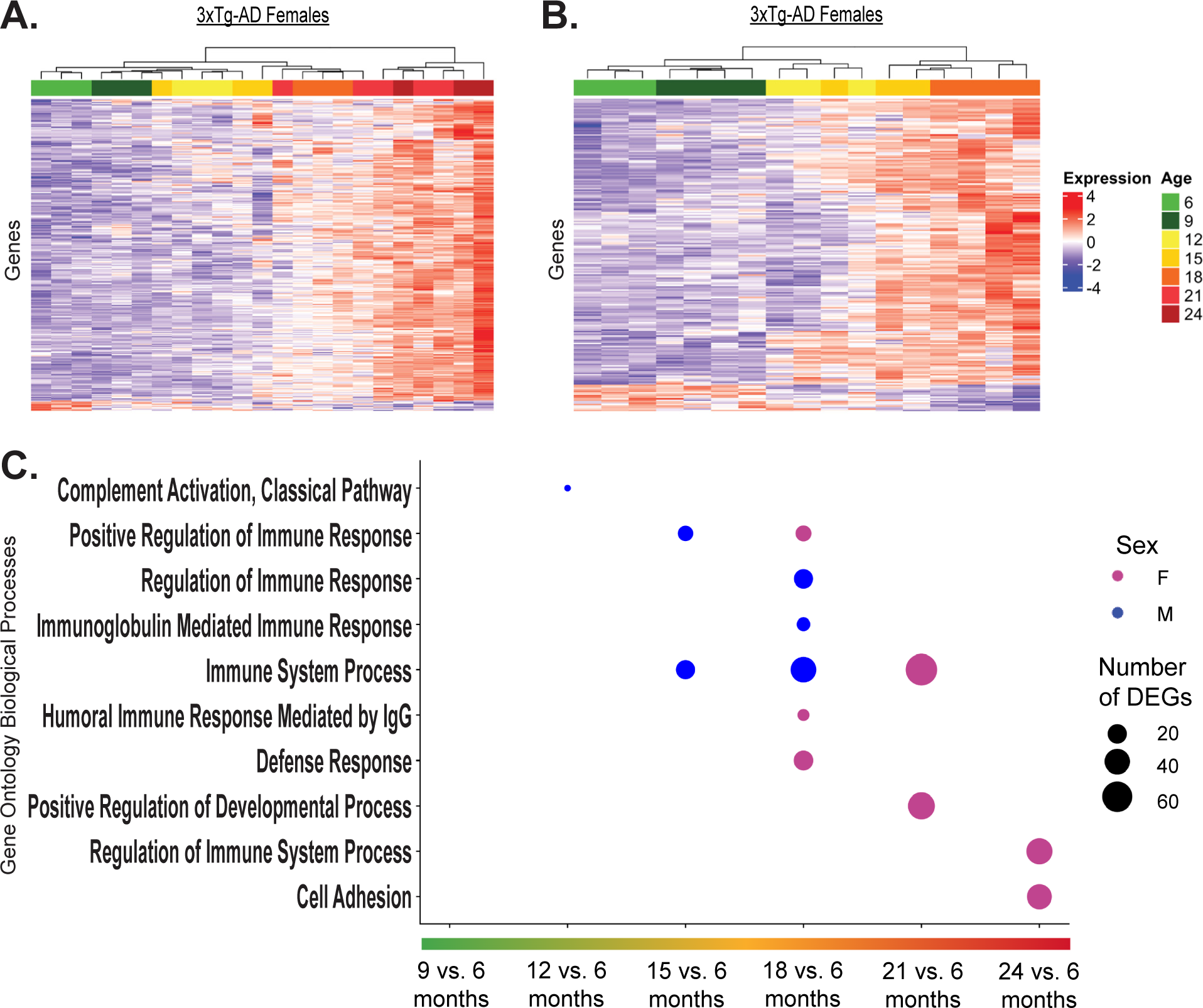
3xTg-AD males exhibit accelerated age-dependent upregulation of immune-related genes in the brain. Total RNA was isolated from the bulk brain tissue of female and male, 6–24-month-old (oldest age dependent upon survival), 3xTg-AD mice and sequenced. Longitudinal differential expression analyses were performed, measuring differences in gene expression of subsequent ages compared to a baseline of 6-months-old (n=3-4/sex+age) with a negative binomial generalized linear model. (A) Heatmap of genes with differential expression in 3xTg-AD females over the course of disease progression (padj < 0.05). Each column represents an individual animal, and age at time of collection is indicated by color. Samples range from 6-24 months of age. (B) Heatmap of genes with differential expression in 3xTg-AD males over the course of disease progression (padj < 0.05). Each column represents an individual animal, and age at time of collection is indicated by color. Samples range from 6-18 months of age. (C) Overrepresentation analysis of significant DEGs between various ages and a 6-month-old baseline. Affinity propagated, overrepresented Gene Ontology biological processes are plotted for females in pink and males in blue. Data were analyzed by one-tailed Fisher’s Exact test. P-values were adjusted for multiple comparisons via Benjamini-Hochberg correction. Number of DEGs indicates number of significant differentially expressed genes from dataset driving enrichment of biological processes.

To understand the foremost biological implications of these molecular changes, we performed overrepresentation analyses on each set of differentially expressed genes, revealing the most enriched Gene Ontology (GO) biological processes over time (Fig. 4c). Additionally, we performed Enrichr pathway analyses on each set of differentially expressed genes, to establish altered pathways over time (Table S3, Table S4). Our analyses demonstrated that for both 3xTg-AD females and males, the majority of enriched biological processes were related to immune regulation; however, the initial age at which significant dysregulation of these processes became evident differed between sexes (Fig. 4c). Beginning at 12-months-old, 3xTg-AD males had enriched complement system activation gene expression in their brains, relative to their 6-month-old baseline. Additional immune-related biological processes were enriched at 15 and 18 months of age, such as immunoglobulin (Ig)-mediated immune response, which included greater numbers of total differentially expressed genes. Contrary to the dysregulation in males that started at 12 months, significantly enriched biological processes were not evident in 3xTg-AD female mice until 18 months of age, relative to their 6-month-old baseline, including humoral immune response mediated by IgG and defense response. Additional immune-related biological processes were further enriched at 21- and 24-months-old. Corresponding analyses were conducted in female and male control mice, revealing minimal molecular changes spanning the ages of 6- to 21-months-old (Sup Fig. 3). These results indicate that the brains of 3xTg-AD mice exhibit highly modulated transcriptomic profiles that worsened with age and can be largely characterized by an upregulation of immune-related processes. Further, 3xTg-AD male mice exhibit accelerated upregulation of immune-related gene expression over the course of disease progression, presenting with significant molecular dysfunction 6-months prior to 3xTg-AD females.

### Aged 3xTg-AD mice and Alzheimer’s patient brains exhibit sex-specific molecular profiles

To determine how the transcriptomic profiles of aged female and male 3xTg-AD brains directly differ from non-diseased controls, we conducted differential expression analyses in 15 and 18-month-old mice (n=6 mice of each sex and strain) (Sup Fig. 4). We found that the resulting statistically significant DEGs could be broken down into three categories: (1) genes that were exclusively differentially expressed in 3xTg-AD females compared to controls, (2) genes that were differentially expressed in both 3xTg-AD females and males compared to controls, or (3) genes that were exclusively differentially expressed in 3xTg-AD males compared to controls. To characterize these sex-specific and overlapping sets of genes, we performed overrepresentation analyses, revealing the most enriched Gene Ontology biological processes in each category (Fig. 5a). Female-specific DEGs were primarily associated with synaptic function, overlapping DEGs were most associated with immune response, and male-specific DEGs were largely associated with additional immune-related processes. To understand further the specific biological pathways of these molecular changes, we performed IPA Core Analysis on significant DEGs. The top 12 altered canonical pathways are plotted for females (Fig. 5c) and males (Fig. 5e). Our analyses indicate that many immune-related pathways, including pathogen-induced cytokine storm signaling and phagosome formation, were dysregulated in 3xTg-AD male mice. In contrast, in 3xTg-AD female mice, various neural signaling pathways were dysregulated including serotonin receptor signaling, dopamine feedback in cAMP signaling, and GPCR signaling. Interestingly, the complement system was altered in both female and male 3xTg-AD mice, with the highest ratio of differentially expressed genes.

**Figure 5.**
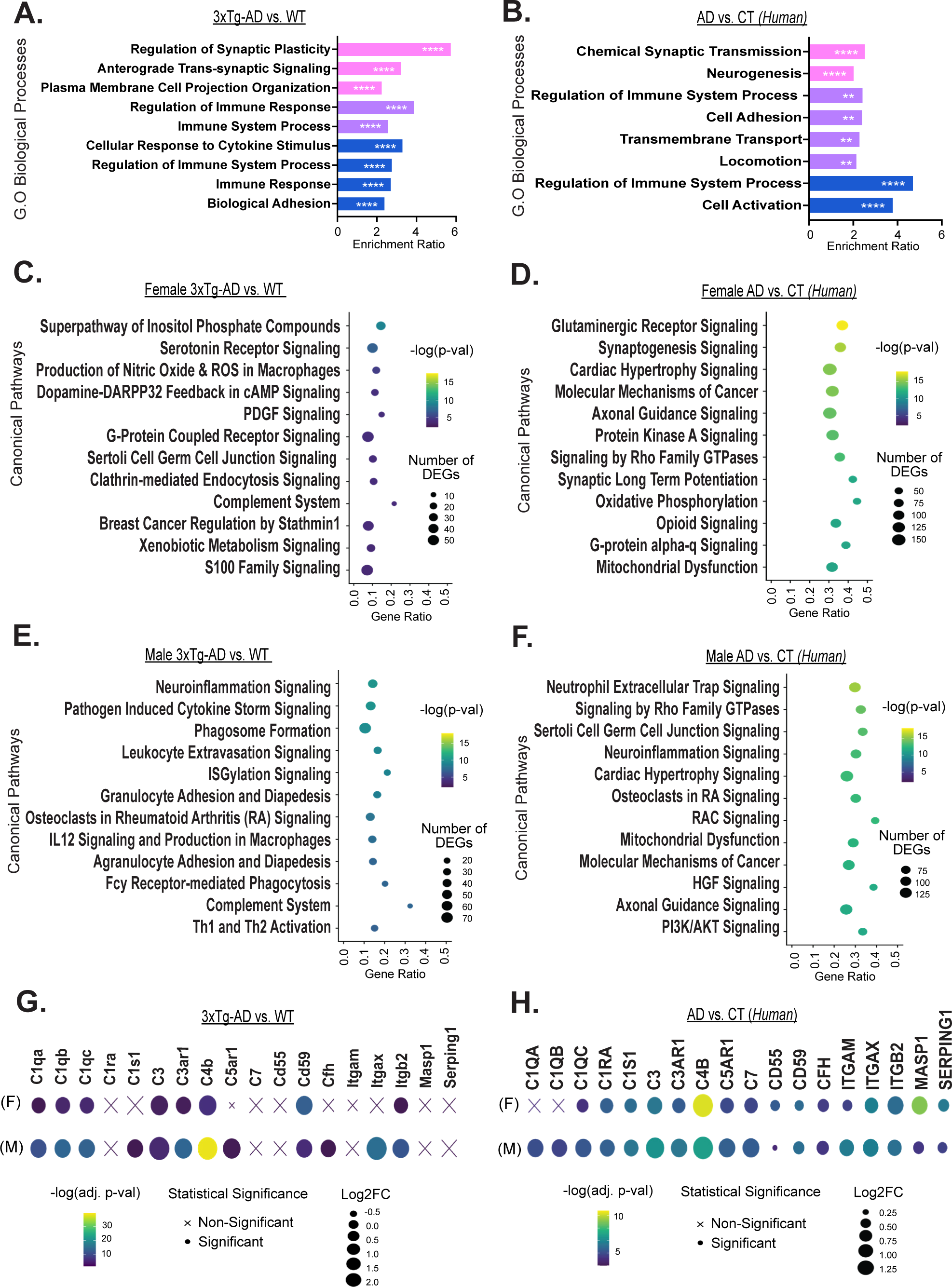
Both aged 3xTg-AD mice and clinical Alzheimer’s disease brains exhibit sex-specific molecular profiles. (A) Total RNA was isolated from the bulk brain tissue of 15–18-month-old, female and male, 3xTg-AD and WT mice, and sequenced. Differential expression analyses were performed between female 3xTg-AD and WT mice (n=6 3xTg-AD, 6 WT) as well as between male 3xTg-AD and WT mice (n=6 3xTg-AD, 6 WT) with a negative binomial generalized linear model. Overrepresentation analysis (ORA) of sex-specific and sex- overlapping significant DEGs (padj < 0.05) in 3xTg-AD mice vs. WT is depicted. Enrichment ratios of significant (FDR < 0.05), affinity propagated, overrepresented Gene Ontology (GO) biological processes, are plotted for female-specific genes in pink, sex- overlapping genes in purple, and male-specific genes in blue. Data were analyzed by one-tailed Fisher’s Exact test. P-values were adjusted for multiple comparisons via Benjamini-Hochberg correction. Significance is indicated with asterisks (** = p≤0.01, **** = p≤0.0001). (B) DEGs in the parahippocampal gyrus (PHG) of females with Alzheimer’s disease (AD) vs. controls (CT) (n=103 AD, 39 CT) as well as males with Alzheimer’s disease vs. controls (n=52 AD, 32 CT) were analyzed. ORA of significant DEGs (padj < 0.001) in individuals with Alzheimer’s compared to controls is depicted. Further details described previously. Top 12 significantly altered canonical pathways in (C) the brains of female 15–18-month-old 3xTg-AD vs. WT, (D) the PHG of females with Alzheimer’s vs. controls, (E) the brains of male 15–18-month-old 3xTg-AD vs. WT, and (F) the PHG of males with Alzheimer’s vs. controls. (C-F) Number of DEGs indicates sum of significant DEGs in dataset driving predicted pathway alteration. Gene ratio indicates number of DEGs out of total genes associated with a specific pathway. Values greater than 1.3 [-log(0.05)] are significant. (G) Complement system gene expression in the brains of 15–18-month-old female and male 3xTg-AD vs. WT, and (H) in the PHG of females and males with Alzheimer’s vs. controls. Log2FC = Log2 fold change.

To validate our transcriptomic findings’ relevance and translational impact in 3xTg-AD mice, we sought to determine how and if the observed sex-specific molecular profiles in our animal model compared to Alzheimer’s disease-related changes in human brains. We examined DEGs in the parahippocampal gyrus of males and females with Alzheimer’s disease compared to controls, obtained from the Mount Sinai Brain Bank study [43]. Again, within the total population of highly significant DEGs, some genes were specifically altered in one sex, while others were differentially expressed in both sexes.

When performing overrepresentation analyses on sex-specific and overlapping sets of genes, the same molecular signatures found in 3xTg-AD mice were observed, as female-specific DEGs were most involved in synaptic function, overlapping DEGs were most associated with immune system process and male-specific DEGs were further associated with the immune system (Fig. 5b). Similarly, IPA Core Analysis revealed altered glutaminergic receptor signaling, synaptogenesis signaling, and axonal guidance signaling pathways in female Alzheimer’s brains (Fig. 5d), while pathways like neutrophil extracellular trap signaling and neuroinflammation signaling were altered in male Alzheimer’s brains (Fig. 5f). These results indicate that both female 3xTg-AD mice and females with Alzheimer’s exhibit a sex-specific molecular phenotype related to neuronal function, while male 3xTg-AD mice and males with Alzheimer’s exhibit an exacerbated neuroinflammatory molecular phenotype.

### Overexpression of complement system genes found in 3xTg-AD mouse and Alzheimer’s patient brains

To dig deeper into the involvement of the complement system in Alzheimer’s disease, we examined the expression of specific complement-related genes. In the brains of 3xTg-AD mice, upregulation of complement-related DEGs including *C1q* and *C4b* indicated that the classical complement pathway was activated (Fig. 5g). All complement-related DEGs had greater log-fold change and more statistical significance in 3xTg-AD males compared to females, and select genes including *C1s1, C5ar1, Cfh,* and *Itgax* only reached statistical significance in males. In the parahippocampal gyrus of humans with Alzheimer’s disease, activation of the classical complement pathway was also revealed; however, further sex differences were observed as *C1QA, C1QB,* and *C1QC* reached statistical significance in males while only *C1QC* reached statistical significance in females (Fig. 5h). Similar to the observations made in 3xTg-AD mice, the majority of complement-related DEGs had greater log-fold change and more statistical significance in males compared to females. However, a few genes were more enriched in females, including *CD55* and *SERPING1*, which both function as complement inhibitors. Another gene with stronger differential expression in females was *MASP1*, activated by the lectin complement pathway which is initiated by the binding of mannose-binding lectin to bacterial surfaces with mannose-containing polysaccharides [44]. This likely also explains the greater statistical significance of *C4B* in females relative to males, as C4 is involved in classical and lectin activation pathways. Overall, our results further substantiate the involvement of complement system activation in Alzheimer’s disease and suggest that sex likely impacts the severity of complement overactivation, with males exhibiting increased upregulation of most complement DEGs, and reduced expression of complement inhibitors, relative to females.

### Alzheimer’s-associated immune cascade resembles downstream effects of LPS stimulation

Based on the abnormal gene expression patterns in female and male Alzheimer’s brains, we used IPA to predict which upstream regulators are likely driving Alzheimer’s-associated differential expression and pathogenesis (Table S5). The top 5 predicted upstream regulators are plotted for females (Fig. 6a,b) and males (Fig. 6c,d). In the mouse and human datasets, we see *MAPT* and *APP* as top predicted upstream regulators, which is to be expected given the animal model and disease pathology.

**Figure 6.**
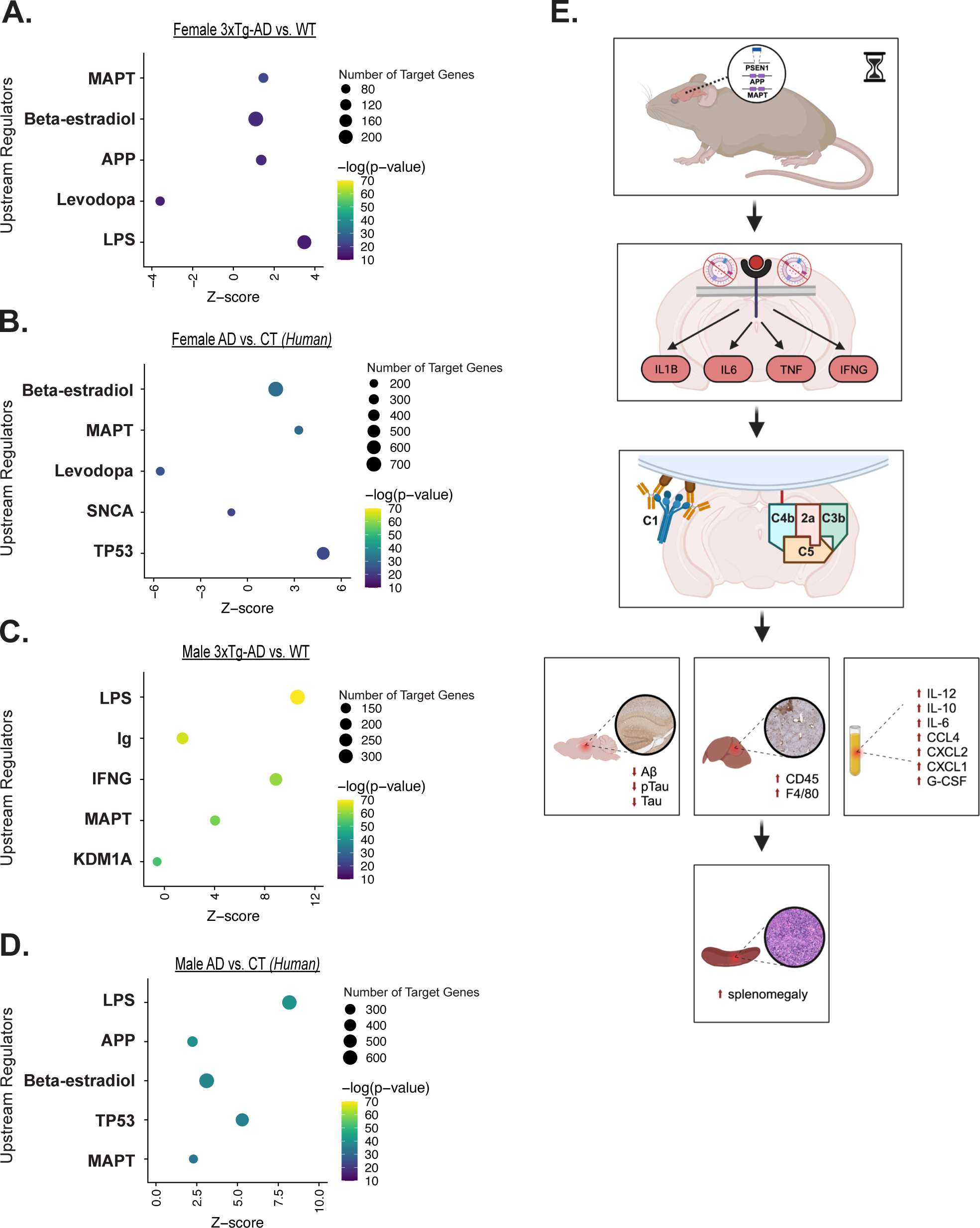
Alzheimer’s disease-associated immune cascade, exacerbated in males, resembles downstream effects of lipopolysaccharide stimulation in mice. (A) Top 5 predicted upstream regulators of DEGs in female 3xTg-AD mouse brains compared to WT. (B) Top 5 predicted upstream regulators of DEGs in female Alzheimer’s disease (AD) parahippocampal gyrus compared to controls (CT). (C) Top 5 predicted upstream regulators of DEGs in male 3xTg-AD mouse brains compared to WT. (D) Top 5 predicted upstream regulators of DEGs in male Alzheimer’s disease parahippocampal gyrus compared to controls. (E) Proposed model of disease pathogenesis in 3xTg-AD males. Overexpressed transgenes are translated in the central nervous system. Aging and disease progression triggers inflammatory response, mimicking effects of LPS stimulation, despite absence of infection. Complement system is activated and associated genes are upregulated. Aβ plaque and NFT formation is prevented. Liver is chronically inflamed, resulting in the development of splenomegaly. Inflammatory cytokines circulate in the bloodstream. *Created with BioRender.com*

However, a striking stand out is that lipopolysaccharide (LPS), an outer-membrane component of gram-negative bacteria and an intense stimulator of the innate immune system, is the number 1 top predicted upstream regulator of gene expression changes in mouse and human male brains. Extensive serology and parasitology analyses have confirmed the absence of pathogens in our utilized animal facility throughout the course of this study (Table S1). Hence, we propose a model in which the Alzheimer’s-associated immune response, clearly exacerbated in males, resembles the downstream effects of LPS stimulation (Fig. 6e). Based on our findings, we hypothesize that mutations in *APP, MAPT,* and *PSEN1* genes yield a compounding effect with age, triggering biological changes which initiate an inflammatory response that mimics the effects of LPS stimulation, despite the absence of infection. Neural activation of additional inflammatory factors, and well-known drivers of inflammaging, including IL1B, IL6, TNF, and IFNG, were also predicted. This process prompts the sustained upregulation of complement system genes, possibly contributing to observed systemic inflammation, hepatic inflammation, and splenomegaly, driving increased mortality. Conversely, chronic inflammatory forces are likely preventing the aggregation of AD-associated proteins in the brains of 3xTg-AD males.

## Discussion

It remains unclear how and when sex differences in biological aging emerge and contribute to Alzheimer’s disease pathogenesis. We hypothesized that aging differentially affects biological processes in males and females, resulting in distinct trajectories of disease. In this study, we first reproduced and then expanded upon previously reported findings in the 3xTg-AD mouse model. Throughout the course of our thirty-month investigation, we recorded all deaths, revealing significantly greater longevity in 3xTg-AD females compared to males (Fig. 1a). We also quantified Alzheimer’s related neuropathology in the brains of female and male 3xTg-AD mice across adulthood (Fig. 2), providing additional evidence that 3xTg-AD females exhibit progressive Alzheimer’s hallmark pathologies. In contrast, 3xTg-AD males exhibit no Aβ plaque pathology and very minimal tangle pathology, if any, in old age. Additionally, we showed that severe splenomegaly was present in aged 3xTg-AD males and associated with abnormal histology (Fig. 3a,b). Together, these results align with published literature [28, 29, 31-33, 45, 46] and reinforce the translatability of the 3xTg-AD model.

To build upon this foundational evidence, we conducted a longitudinal analysis of liver inflammation in female and male 3xTg-AD and control mice. While others have reported hepatomegaly in the 3xTg-AD model [45-47], we are the first, to our knowledge, to explore hepatocellular expression of CD45 and F4/80 across sex, age, and disease status. We discovered that male 3xTg-AD mice exhibit significant chronic hepatic inflammation throughout adulthood, while females exhibit milder hepatic inflammation in old age (Fig. 3c-f). These results help to elucidate the driver of splenomegaly in male 3xTg-AD mice and raise questions regarding the involvement of the liver and liver-brain axis dysfunction in Alzheimer’s disease. Evidence has pointed to the liver as a target organ in Alzheimer’s due to its role in regulating metabolism and supporting the immune system [48-50]. Researchers have proposed that impaired liver function due to chronic liver diseases and inflammation leads to an imbalance in peripheral Aβ clearance, exacerbating amyloid burden and Alzheimer’s pathology [51, 52]. However, in our study, male 3xTg-AD mice with severe hepatic inflammation did not exhibit plaque pathology. This implies that the liver likely has a broader role outside of amyloid clearance alone in its contributions to Alzheimer’s pathogenesis.

Heightened levels of CD45+ and F4/80+ cells have been described in the livers of 24-month-old male C57BL/6 mice [53] and Fischer rats [54]. While we did not observe evidence of increasing CD45 or F4/80 staining in our controls at up to 18-months of age (Fig. 3b-e), it is possible that there is a delayed or rapid onset of age-related hepatic inflammation in B6129 mice. Further, the hepatic inflammation observed in male 3xTg-AD mice could be the result of an accelerated aging phenotype in the male Alzheimer’s disease trajectory. While limited studies have investigated the impact of aging on hepatic inflammation and function in humans, strong evidence supports the connection between liver and brain-related maladies, as non-alcoholic fatty liver disease has been linked to an increased risk of dementia [55]. A clinical study of over 450,000 participants [56] revealed that males were more likely to experience liver fibrosis and exhibited higher associations between liver fibrosis and most cognitive performance tests, suggesting that hepatic inflammation has a greater neurological impact in males than in females. However, the precise relationship between liver pathology, neuropathology, and neurodegeneration remains unclear.

To elucidate the temporal relationship between hepatic and neuroinflammation in the 3xTg-AD mouse model, we performed sex-specific longitudinal analyses of age and disease-related neural gene expression changes (Fig. 4). This study is one of the first to delineate sex-specific molecular timelines of disease progression in the 3xTg-AD model, revealing accelerated rates of neuroinflammation in male 3xTg-AD mice through the overrepresentation of immune-related processes beginning at 12-months-old in males and 18-months-old in females. These results were initially surprising, given the escalating presence of Alzheimer’s neuropathology in female 3xTg-AD mice from 12-months of age onward, in addition to the commonly accepted paradigm that females develop stronger immune responses than males. However, age is a key variable that influences the strength and effectiveness of immune responses. Male and female immune responses change greatly over the life course [20]. Post-puberty and throughout the majority of adulthood, females exhibit heightened innate and adaptive immunity compared to males, resulting in 80% of autoimmune disease occurrence and greater vaccine efficacy in women. However, in old age, inflammation associated with innate immunity is heightened in males compared to females. The rate of “inflammaging” is also greater in men [21-23], driving a gap in male and female life expectancies. This process likely plays a key role in the pathogenesis of age-related diseases, including Alzheimer’s [24].

A key contributor to the development and progression of inflammaging is the senescence associated secretory phenotype (SASP) – a damaging byproduct of senescent cells [57], and a potential driver of Alzheimer’s pathogenesis [58]. Cellular senescence is a tumor suppression mechanism which prevents the uncontrolled proliferation of old and injured cells. Despite the protective intent of this mechanism, senescent cells are accompanied by SASP which is comprised of secreted proteases and insoluble proteins, as well as soluble signaling factors such as interleukins, chemokines, and growth factors. Senescence is widely accepted as a major cause of age-related disease. The accumulation of senescent cells drives chronic immune induction, which in turn impairs the clearance of senescent cells, creating a detrimental cycle that fuels inflammaging [57]. We showed that aged 3xTg-AD male mice in particular, exhibit elevated plasma levels of known SASP factors [59, 60], including IL-12, IL-10, IL-6, CXCL2, CXCL1, and G-CSF (Sup Fig. 1). Together, our data in the liver, spleen, plasma, and brain clearly implicate chronic immune upregulation in Alzheimer’s disease-associated mortality. Thus, we predict that biological aging and consequential age-associated inflammation are the central driver behind the sex differences observed in 3xTg-AD mice.

To further elucidate the impact of disease progression on the transcriptomic profiles of aged 3xTg-AD brains, we performed differential expression analysis with direct comparison to controls, which revealed a female-specific primary disease phenotype associated with impaired synaptic function and neurotransmitter signaling pathways (Fig. 5a,c). These results imply that 3xTg-AD females experience more neurodegeneration than males; therefore, the neuroinflammation observed in female and male 3xTg-AD brains may be the consequence of distinct forces. In Alzheimer’s disease, neuroinflammation is primarily thought to be a consequence of a series of damage signals including trauma, oxidation, oligomers of Aβ and tau, and more [61]. Immune processes are triggered to maintain homeostasis and protect the neural environment. Eventually, however, neuroinflammation becomes chronic, initiating a cyclic cascade of glial priming, proinflammatory factor release, and neuronal damage [62]. Neuropathological and molecular changes associated with neuroinflammation in 3xTg-AD females align with this described pathogenic mechanism. Progressive plaque and tangle pathology were first observed in 3xTg-AD females at 12 months and 15 months of age respectively, subsequently followed by escalating neuroinflammation between 18 months and 24 months of age (Fig. 2a-d, Fig. 4c). However, neuroinflammation in 3xTg-AD males does not follow this timeline, suggesting that an earlier trigger kicks off an unrelenting immune cascade, driven by earlier disease abnormalities as opposed to later stage disease pathology.

One key similarity between the sexes, and one of the most clearly perturbed biological pathways in our study, was an overactive complement system, a vital element of the immune system that enhances the opsonization of pathogens and initiates a series of inflammatory responses to help fight infection [44]. The primary action of the complement system is to enable the uptake and destruction of pathogens by phagocytic cells, which are microglia in the brain. In aged 3xTg-AD mice, the complement system had the greatest ratio of DEGs among top altered canonical pathways in both male and female brains (Fig. 5g). Upregulation of *C1q*, which is known to bind to Aβ peptides [63], NFT filaments [64], apoptotic cells and extracellular debris [65], indicated that the classical activation pathway was activated. Interestingly, 3xTg-AD males exhibited more extensive complement system overactivation than females, as differentially expressed complement genes were more statistically significant with greater log-fold changes.

Strong evidence has implicated the complement system in the pathogenesis of Alzheimer’s disease [66-70], but questions remain regarding the impact of sex on complement activation and microglial phagocytosis. Several studies point to more potent and plentiful complement factors, and higher rates of microglial phagocytosis in males [71-74], while others find the opposite sex-difference in females [75-77]. Similar to other immune system studies previously described, age is likely a key variable between these experiments, which is especially difficult to capture in *in-vitro* studies, and strongly influences sex-specific results. Further, studies manipulating the complement system have yielded mixed results. The deficiency of complement factors such as C1q, C3, C3ar1, C5aR, and Cd59, has exacerbated neuropathology in some studies [78-80] and reduced neuropathology in others [81-83]. Additionally, multiple studies have shown that complement factor inhibition ameliorates synapse and neuron loss [84-87]. Determining how and when to modulate the complement cascade to prevent abnormal activation will be crucial, and more longitudinal studies will aid in identifying the appropriate and helpful window of treatment. Our transcriptomic results suggest that targeting C1q could serve as a targeted solution to both harness protective actions and dampen destructive actions of the complement system.

To explore further the primary drivers of differential gene expression in aged 3xTg-AD brains, we performed upstream regulator analyses. Female and male 3xTg-AD neural transcriptomes yielded distinct upstream regulator predictions. Differential gene expression was predicted to be most influenced by LPS, immunoglobulin, and interferon-gamma activation in male 3xTg-AD brains (Fig. 6c), as opposed to *MAPT*, beta-estradiol, and *APP* in female 3xTg-AD brains (Fig. 6a). These results further reinforce that male 3xTg-AD mice experience an inflammatory manifestation of disease. Further comparison between the strength of upstream regulator predictions in males and females revealed that the other proinflammatory factors [88-90] most associated with inflammaging – IL1B, IL6, and TNFα – all have statistically greater predictions of activation in male 3xTg-AD brains, with z-scores three to fifteen-fold larger than that of females. We were surprised to discover that LPS, an outer-membrane component of gram-negative bacteria and an intense stimulator of the innate immune system, is the leading top predicted upstream regulator of gene expression changes in the male 3xTg-AD brain. Serum screening in sentinel mice confirmed the absence of any known pathogens (Table S1). Therefore, an endogenous abnormality is triggering a pathogen-like response in 3xTg-AD males.

To broaden the translational impact of our work, we sought to verify the results obtained from our animal model in clinical data. We analyzed DEG data generated by The Mount Sinai/JJ Peters VA Medical Center NIH Brain and Tissue Repository, comparing the parahippocampal gyri of female and male Alzheimer’s brains to age-matched controls [43]. We found overwhelming similarities between the mouse and human data. We again observed a female-specific primary disease phenotype associated with impaired synaptic function, implying advanced neurodegeneration in the female Alzheimer’s parahippocampal gyrus (Fig. 5b). Additionally, the top altered canonical pathway in the male Alzheimer’s parahippocampal gyrus was neutrophil extracellular trap signaling, an immune process activated by pathogens, antibodies, cytokines, and other physiological stimuli (Fig. 5f). In both the 3xTg-AD mouse model, and in humans with Alzheimer’s, we see sex-specific disease phenotypes primarily driven by compromised neuronal function in females and aggravated immune response in males. Further, detailed gene expression analyses revealed significant upregulation of complement system genes in male and female Alzheimer’s parahippocampal gyri (Fig. 5h). Both animal and human results suggest that sex may impact the extent of complement system activation, as males exhibited a heightened upregulation of complement system gene expression in the brain. Furthermore, the leading predicted upstream regulator of differential gene expression in the male Alzheimer’s parahippocampal gyrus was LPS, precisely matching our observations in the male 3xTg-AD brain.

The infectious hypothesis proposes that a pathogen is the primary driver of Alzheimer’s disease [91, 92]. Studies have linked oral bacteria [93], the gut microbiome [94], herpesviruses [95], and other environmental pathogens with an increased risk of dementia. Theories regarding the involvement of Alzheimer’s neuropathology in the infectious hypothesis differ. Plaques and tangles are posed to either associate with pathogens directly or develop as a result of infection-driven inflammation. Given that LPS was predicted as the leading upstream regulator in males of both species, our results suggest that females are less impacted by pathogens or an endogenous pathogen-like trigger, and that males are more susceptible to an infectious mode of Alzheimer’s initiation. Hence, we suggest that the male-biased immune manifestations of Alzheimer’s disease resemble the downstream effects of LPS stimulation (Fig. 6e). We hypothesize that the mutations in *APP, MAPT,* and *PSEN1* genes, yield a compounding effect with age. The resulting biological changes initiate a neuroinflammatory response that mimics the effects of LPS stimulation, likely characterized by microglial activation. Pro-inflammatory factors, and well-known drivers of inflammaging, are activated including IL1B, IL6, TNFα, and IFNγ, in addition to the complement system. We hypothesize that persistent complement-mediated microglial phagocytosis clears and prevents the formation of plaques and tangles in the male 3xTg-AD brain, while the chronic inflammatory environment results in liver inflammation, spleen enlargement, and circulating inflammatory proteins in the bloodstream, driving increased mortality.

This hypothesis is substantiated by published literature revealing sex differences in microglial signature and response to complement receptor agonism. Hanamsager et al. [96] identified a microglia-specific gene expression program in mice which enabled the generation of a microglial developmental index. This work showed that in response to LPS injection, male C57BL/6 microglia elicit an aged phenotype characterized by heightened upregulation of immune gene expression relative to female derived microglia. Additionally, healthy human male brains were found to exhibit “older” microglial signatures than age-matched female brains. Brains from individuals with Alzheimer’s disease were also found to exhibit “older” microglial signatures than age-matched controls. While sex-specific analyses were not conducted with Alzheimer’s data in this study, our results support the prediction that microglia from male Alzheimer’s brains would exhibit the “oldest” reactive microglial signature.

Further, Stephen et al. [97] examined how sex and *APOE* genotype affect microglial interactions with plaques in the EFAD Alzheimer’s mouse model. They discovered that microglial coverage of amyloid plaques is highest in male *APOE3* mice and reduced by female sex and *APOE4* genotype. Thus, female sex contributes to impaired microglial protective action in the Alzheimer’s brain, supporting our hypothesis that males exhibit lower Aβ plaque burden as a result of exacerbated immune response. Gaamouch et al. [80] treated 5xFAD mice with a VGF-derived peptide which activates complement C3a receptor-1 (C3aR1), predominantly expressed in the brain on microglia. Interestingly, treatment significantly reduced Aβ plaque load in males but not in females. This work provides additional evidence that the complement system regulates Alzheimer’s neuropathology and suggests that either the male immune system is more effective at clearing and preventing protein aggregation, or that advanced pathological burden in the female brain reaches a point of no return. Lastly, our results also align with Kapadia et al. [98] who showed that chronic immunosuppression via cyclophosphamide treatment prevented hepatosplenomegaly in 3xTg-AD mice. The group has also reported that 3xTg-AD males undergo a sex-specific systemic autoimmune response [45, 46]. While we did not examine markers of autoimmunity in our study, it is possible that an autoimmune reaction is contributing to the male inflammatory disease phenotype observed in our study. Our analyses of clinical Alzheimer’s disease data confirm that widespread inflammation in 3xTg-AD males is not simply an artifact of the animal model, and that further investigation into the mechanisms behind sex-specific immune responses in Alzheimer’s disease is warranted.

## Conclusions

In conclusion, our data demonstrate that chronic inflammation and complement activation are associated with increased mortality, revealing that age-related changes in immune response act as a primary driver of sex differences in Alzheimer’s disease trajectories. Current “disease-modifying” FDA-approved therapies for Alzheimer’s are anti-amyloid antibody drugs, which successfully eliminate amyloid fibrils from the brain, but have minimally effective benefits in regard to disease progression and cognitive protection. Our study shows that destructive neuro- and systemic inflammation occurs without the presence of Aβ plaque pathology in the male 3xTg-AD mouse model, raising questions regarding the true impact of the protein in the progression of disease. Further, we demonstrate that males and females manifest distinct timelines of Alzheimer’s disease progression and will require a personalized medicine approach to prevent and treat dementia most effectively. The data presented in this study provides evidence that aging affects the immune system of males and females differently. Sex-differences in inflammaging must be further studied in order to develop effective therapeutics that incorporate patient demographics and evolve past a one-size-fits-all approach to treating patients with Alzheimer’s disease.

## Abbreviations

Aβ: Amyloid Beta
AD: Alzheimer’s disease
ANOVA: Analysis of Variance
CCL: CC chemokine ligand
CD45: Leukocyte common antigen
CT: Controls (human)
CXCL: CXC chemokine ligand
DEGs: Differentially Expressed Genes
EDTA: Edetic Acid
G-CSF: Granulocyte-Colony Stimulating Factor
GO: Gene Ontology
Ig: Immunoglobulin
IL: Interleukin
IPA: Ingenuity Pathway Analysis
LPS: Lipopolysaccharide
MSBB: Mount Sinai Brain Bank
NFTs: Neurofibrillary Tangles
ORA: Overrepresentation Analysis
PHG: Parahippocampal Gyrus
RNA-Seq: RNA Sequencing
SASP: Senescence Associated Secretory Phenotype
WT: Wildtype (B6129 mice)

## Acknowledgements

RNA sequencing was carried out in the Genomics and Molecular Biology Shared Resource (GMBSR) at Dartmouth which is supported by NCI Cancer Center Support Grant 5P30CA023108 and NIH S10 (1S10OD030242) awards. We especially thank Dr. Fred Kolling, PhD for his guidance and experimental support. The skills and tools necessary to conduct RNA sequencing data analyses were acquired through the Data Analytic Core at Dartmouth. We thank Dr. Shannon Soucy, PhD for her instruction, advice, and technical support. The Center for Comparative Medicine and Research (CCMR) at Dartmouth maintained animal cages, monitored animal safety, provided animal handling trainings, and performed retro-orbital bleeds. We thank Kathryn Bennett and Stephen Gunn for their services and care. The Pathology Shared Resource at Dartmouth performed tissue processing, antibody staining, and slide imaging for all immunohistochemistry experiments. We thank Scott Palisoul for his technical services, antibody selection guidance, and protocol optimization feedback. Luminex experiments were carried out in DartLab, the Immune Monitoring and Flow Cytometry Shared Resource at the Norris Cotton Cancer Center at Dartmouth, with NCI Cancer Center Support Grant 5P30 CA023108-41. We thank Dr. Daniel Mielcarz, PhD and Andrew Calkins for their support and technical services. We thank Drs. Ta Yuan Chang, PhD, Francesca Gilli, PhD, and Lucas Salas, MD, MPH, PhD for their expert guidance and manuscript review.

## Availability of data and materials

The results published here are in part based on data obtained from the AD Knowledge Portal (https://adknowledgeportal.org). Data generation was supported by the following NIH grants: P30AG10161, P30AG72975, R01AG15819, R01AG17917, R01AG036836, U01AG46152, U01AG61356, U01AG046139, P50 AG016574, R01 AG032990, U01AG046139, R01AG018023, U01AG006576, U01AG006786, R01AG025711, R01AG017216, R01AG003949, R01NS080820, U24NS072026, P30AG19610, U01AG046170, RF1AG057440, and U24AG061340, and the Cure PSP, Mayo and Michael J Fox foundations, Arizona Department of Health Services and the Arizona Biomedical Research Commission. We thank the participants of the Religious Order Study and Memory and Aging projects for the generous donation, the Sun Health Research Institute Brain and Body Donation Program, the Mayo Clinic Brain Bank, and the Mount Sinai/JJ Peters VA Medical Center NIH Brain and Tissue Repository. Data and analysis contributing investigators include Nilüfer Ertekin-Taner, Steven Younkin (Mayo Clinic, Jacksonville, FL), Todd Golde (University of Florida), Nathan Price (Institute for Systems Biology), David Bennett, Christopher Gaiteri (Rush University), Philip De Jager (Columbia University), Bin Zhang, Eric Schadt, Michelle Ehrlich, Vahram Haroutunian, Sam Gandy (Icahn School of Medicine at Mount Sinai), Koichi Iijima (National Center for Geriatrics and Gerontology, Japan), Scott Noggle (New York Stem Cell Foundation), Lara Mangravite (Sage Bionetworks).

## Author Information

### Contributions

AB contributed to study design, maintained the animal colony, was responsible for all aspects of data acquisition and analysis, and wrote the manuscript. CD contributed to study design and data acquisition by initiating animal colony breeding, assisting with animal colony maintenance, and assisting with biospecimen collection. AS contributed to data acquisition by assisting with the development and execution of the immunohistochemistry quantification protocol, biospecimen collection and organization, and animal colony maintenance. EP contributed to data acquisition by assisting with biospecimen collection and animal colony maintenance. SW contributed to data acquisition by assisting with immunohistochemistry quantification. VT assisted with review of RNA-Seq data. GZ supervised immunohistochemistry work as well as reviewed and revised the manuscript. AG developed and supervised the study, as well as reviewed and revised the manuscript. All authors approved the final version of the manuscript.

## Ethics declarations

### Ethics approval and consent to participate

All applicable national and institutional guidelines for the care and use of animals were followed.

## Supplementary Information

**Additional File 1: Table S1.** Serology testing agents.

**Additional File 2: Table S2.** Abnormalities observed during animal necropsy.

**Additional File 3: Table S3.** Top altered pathways between ages in 3xTg-AD and WT females.

**Additional File 4: Table S4.** Top altered pathways between ages in 3xTg-AD and WT males.

**Additional File 5: Table S5.** Predicted upstream regulators in female and male 3xTg-AD and AD brains.

**Additional File 6: Supplementary Figures 1-4.**

