## Supplementary Figures 1-4 for "Age, Sex and Alzheimer’s disease: A longitudinal study of 3xTg-AD mice reveals sex-specific disease trajectories and inflammatory responses mirrored in postmortem brains from Alzheimer’s patients"

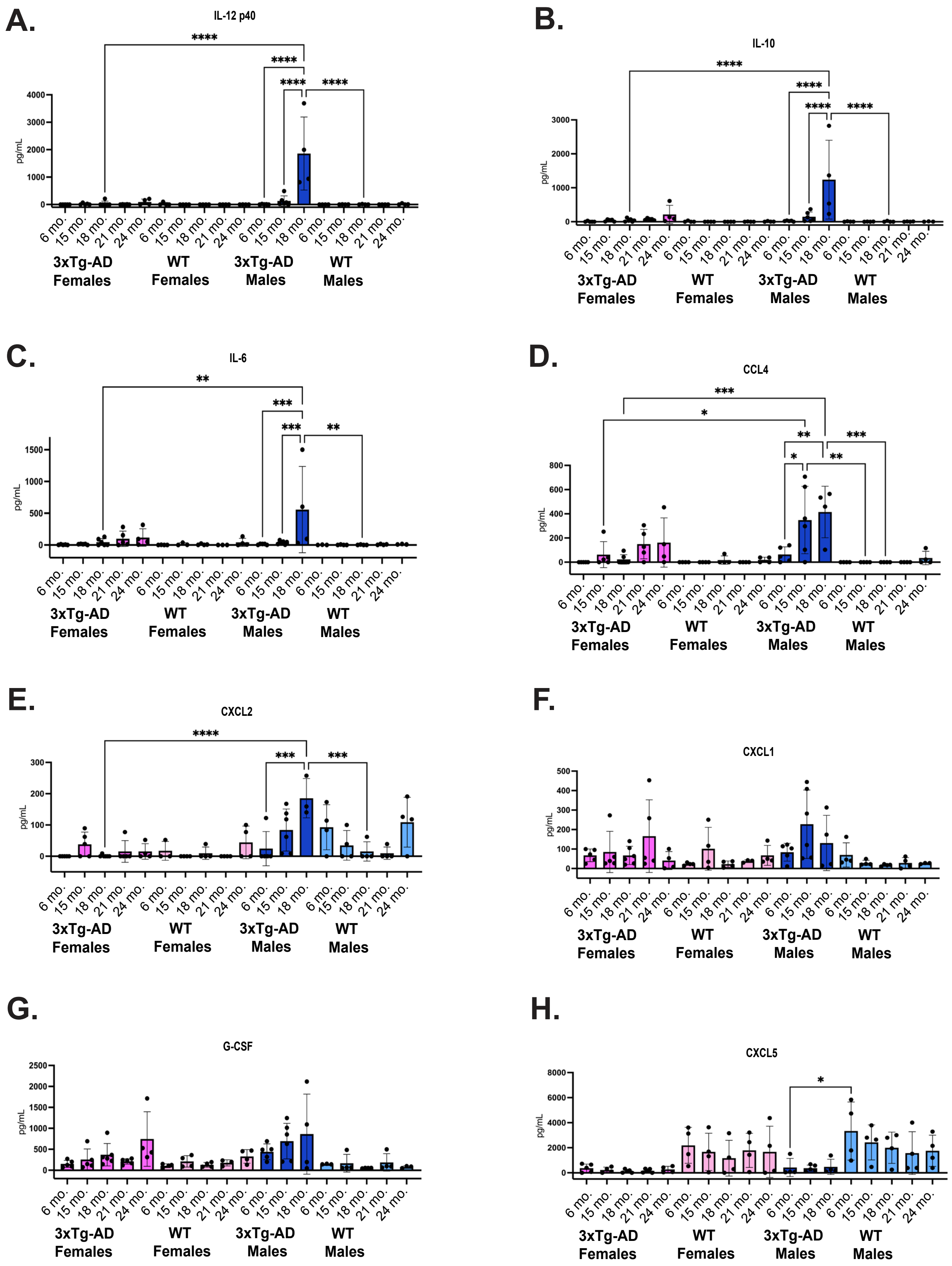
A.
IL-12 p40
B.
IL-10
C.
IL-6
D.
CCL4
E.
CXCL2
F.
CXCL1
G.
G-CSF
H.
CXCL5
pg/mL
6 mo.
15 mo.
18 mo.
21 mo.
24 mo.
3xTg-AD Females
WT Females
3xTg-AD Males
WT Males

I.

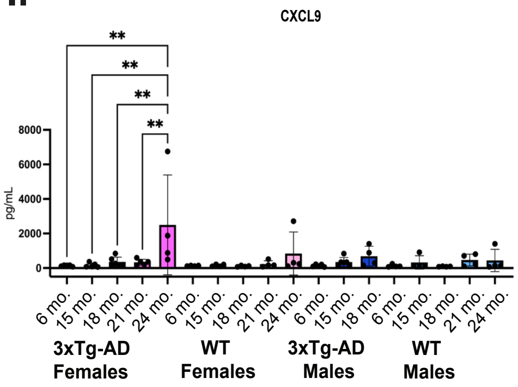


J.

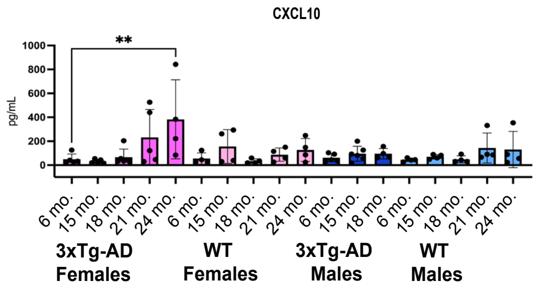


Supplementary Figure 1. **3xTg-AD males exhibit an age-dependent increase in circulating inflammatory proteins.** Cytokine concentrations were measured in plasma from female and male, 3xTg-AD and WT mice, at ages 6-, 15-, 18-, 21-, and 24-months (oldest age dependent upon survival) (n=3-6/sex+age+strain). Observed plasma concentrations of (A) IL-12 p40, (B) IL-10, (C) IL-6, (D) CCL4, (E) CXCL2, (F) CXCL1, (G) G-CSF, (H) CXCL5, (I) CXCL9, and (J) CXCL10. Data points considered out of range below were substituted with a 0, the minimum detectable value. Data were analyzed by one-way ANOVAs with multiple comparisons using GraphPad Prism. P-values less than 0.05 were considered statistically significant. Asterisks indicate significant differences between compared groups (* = $p<0.05$, ** = $p\leq0.01$, *** = $p\leq0.001$, **** = $p\leq0.0001$). Error bars represent mean with SD.

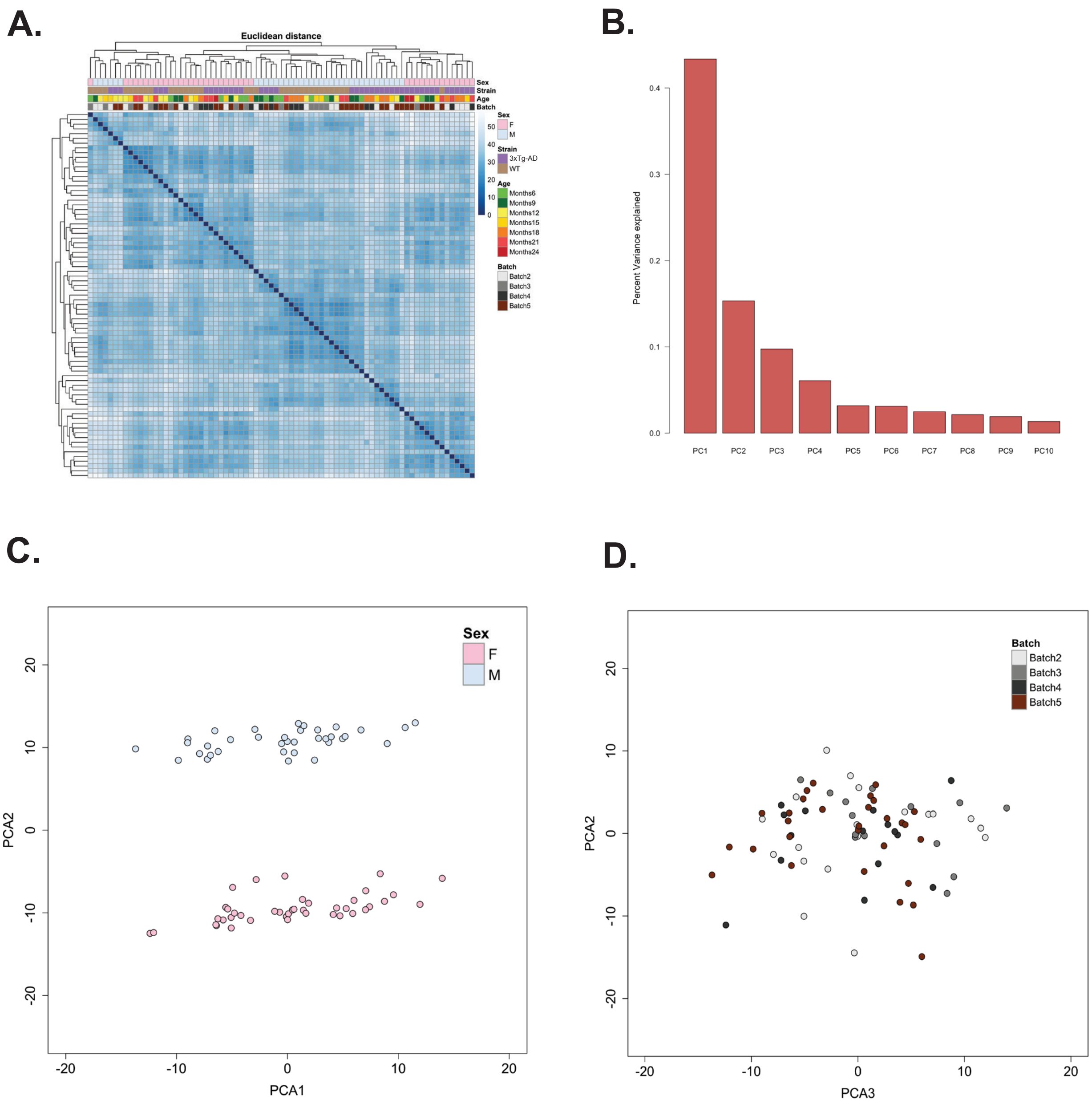


Supplementary Figure 2. **Exploratory analysis of complete RNA sequencing sample pool.** (A) Hierarchical clustering based on Euclidian distance. Euclidian distance values for each sample compared to all other samples are plotted. Sample characteristics including sex, mouse strain, age, and sequencing batch are indicated. (B) Variance of top 10 principal components. (C) Principal component 1 plotted against principal component 2, showing complete sample clustering based on sex. (D) Principal component 3 plotted against principal component 2, showing no strong sample clustering based on sequencing batch.

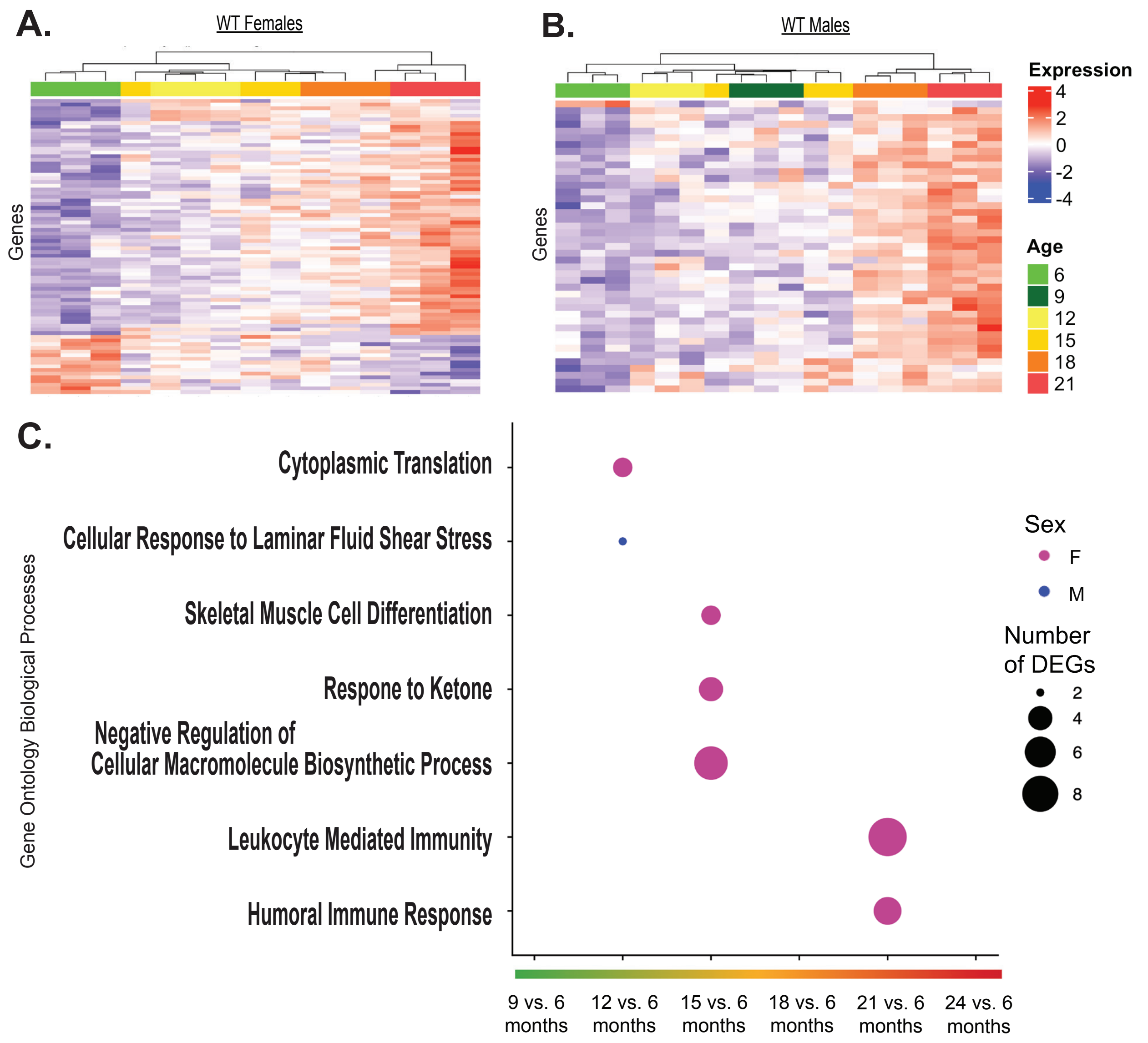


Supplementary Figure 3. **Longitudinal gene expression in control mice.** Total RNA was isolated from the bulk brain tissue of female and male, 6-21 month-old, WT mice and sequenced. Longitudinal differential expression analyses were performed, measuring differences in gene expression of subsequent ages compared to a baseline of 6-months-old (n=3-4/age). (A) Heat-map of genes with differential expression in WT females over the course of disease progression (padj < 0.05). Each column represents an individual animal, and age at time of collection is indicated by color. Samples range from 6-21 months of age. (B) Heatmap of genes with differential expression in WT males over the course of disease progression (padj < 0.05). Each column represents an individual animal, and age at time of collection is indicated by color. Samples range from 6-21 months of age. (C) Overrepresentation analysis of significant DEGs between various ages and a 6-month-old baseline. Affinity propagated, overrepresented GO biological processes are plotted for females in pink and males in blue. Data were analyzed by one-tailed Fisher's Exact test. P-values were adjusted for multiple comparisons via Benjamini-Hochberg correction. Number of DEGs indicates number of significant differentially expressed genes from dataset driving enrichment of biological processes.

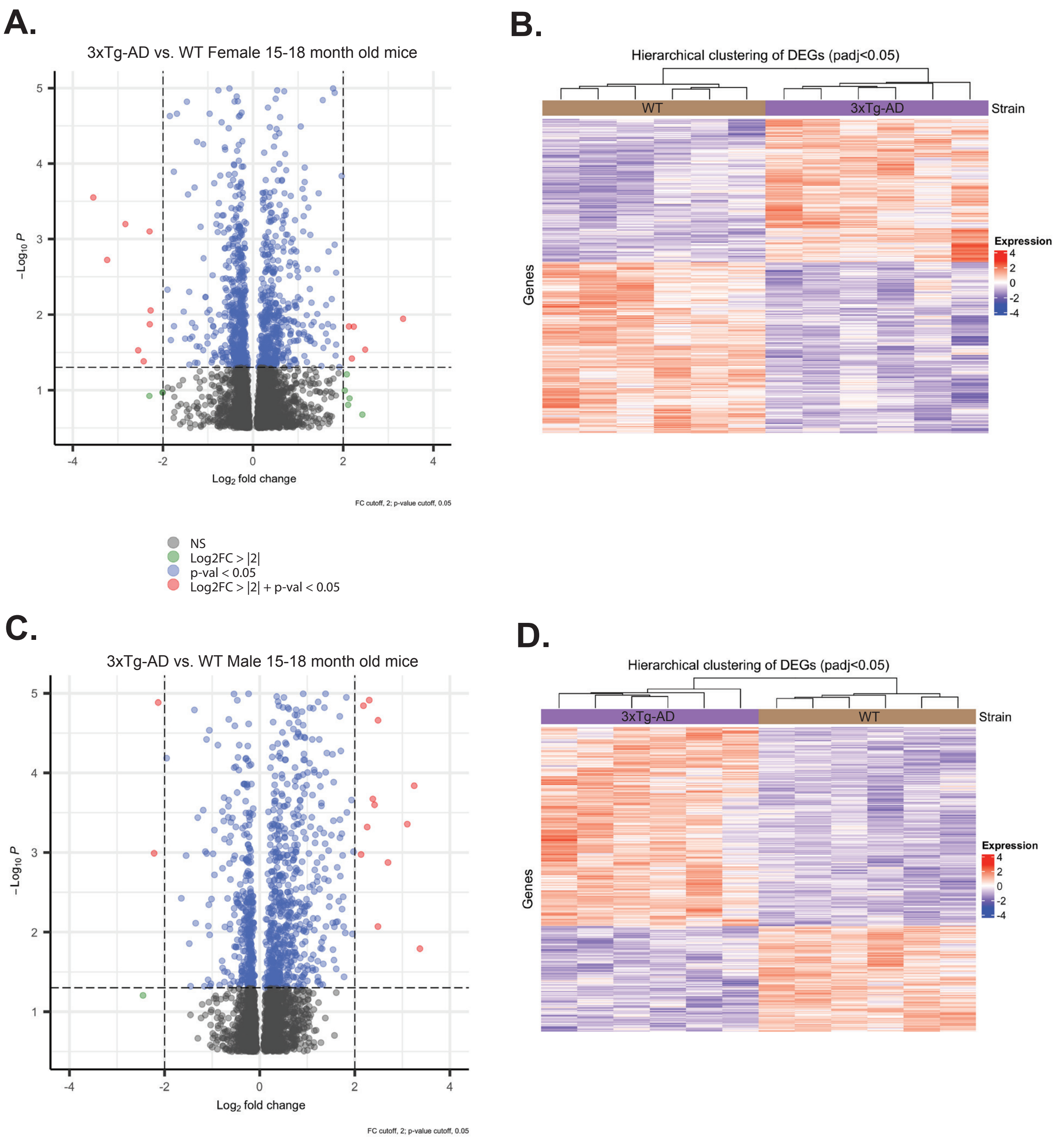


Supplementary Figure 4. **Visualization of differential gene expression in 15-18 month old mice.** Differential expression analyses of RNA from bulk brain tissue were conducted between female and male 3xTg-AD and WT mice at 15-18 months of age. (A) Log2 fold change of each gene plotted against its adjusted p-value. Difference in gene expression between females. (B) Hierarchical clustering of all significant differentially expressed genes (padj<0.05). 3xTg-AD female samples cluster separately from WT female samples. (C) Log2 fold change of each gene plotted against its adjusted p-value. Difference in gene expression between males. (D) Hierarchical clustering of all significant differentially expressed genes (padj<0.05). 3xTg-AD male samples cluster separately from WT male samples.
